# Analysis of a large data set reveals haplotypes carrying putatively recessive lethal alleles with pleiotropic effects on economically important traits in beef cattle

**DOI:** 10.1101/414292

**Authors:** Janez Jenko, Matthew C McClure, Daragh Matthews, Jennifer McClure, Martin Johnsson, Gregor Gorjanc, John M Hickey

## Abstract

**Background:** Deleterious recessive alleles can result in reduced economic performance in livestock in multiple ways in homozygous individuals: from early embryonic death, death soon after birth, to being non-lethal but causing reduced viability. While death is an easy phenotype to score, reduced viability is not as easy to identify. However, it can sometimes be observed as reduced artificial insemination (AI) conception rates, longer calving intervals, or higher hazard for live born animals.

**Methods:** In this paper, we searched for haplotypes carrying putatively recessive lethal alleles in 132,725 genotyped Irish beef cattle from five breeds: Aberdeen Angus, Charolais, Hereford, Limousin, and Simmental. We phased the genotypes in sliding windows along the genome and used five tests to identify haplotypes with absence of or reduced homozygosity. We then corroborated the identified haplotypes with reproduction records, indicating early embryonic death, and postnatal survival records. Finally, we assessed haplotype pleiotropy by estimating substitution effects on national estimates of breeding values for 15 economically important traits in beef production.

**Results:** We found support for three haplotypes with carrying putatively recessive lethal alleles. The haplotypes were located on chromosome 14 in Aberdeen Angus, chromosome 19 in Charolais and chromosome 16 in Simmental. Their population frequencies is 15.2%, 14.4%, and 8.8%, respectively. All of the haplotypes showed pleiotropic effects on economically important traits for beef production. Their allele substitution effects are €3.23, €1.47, and €2.30 for the terminal index and -€3.15, -€0.75, and €1.12 for the replacement index, where one standard deviations are €18.32, €22.54, and €22.33 for terminal index and €29.52, €35.62, and €30.97 for the replacement index. We identified *ZFAT* as the candidate gene for lethality in Aberdeen Angus, several candidate genes for the Simmental haplotype, and no candidate genes for the Charolais haplotype.

**Conclusions:** We analysed genotype, reproduction, survival, and production data to discover haplotypes carrying putatively recessive lethal alleles in Irish beef cattle. We found support for three haplotypes. All three haplotypes have pleiotropic effects on economically important traits in beef production.

## Background

Reproduction inefficiency has a major negative effect on profitability in cattle farming [1–4]. Over the last few decades, reproduction has deteriorated in both dairy and beef cattle in many countries, due to strong selection on production traits and their antagonistic relationship with reproduction [5,6] and probably also due to genetic drift. In recent years, this trend has begun to be readdressed in many countries by emphasising reproduction traits in animal recording and breeding goals [7,8]. Further, the advent of affordable high-density genotyping has enabled genomic selection jointly for production and reproduction traits [9,10] and screening for recessive lethal alleles [11–15].

Recessive lethal alleles are one of the genetic reasons for reproduction inefficiency. They cause early embryonic death when an embryo is homozygous. Such alleles tend to be rare, but if their frequency increases, the number of affected (homozygous) embryos increases and so does the economic loss. Recessive lethals may increase in frequency in a population because of genetic drift, linkage with favourable alleles, or heterozygote advantage. For example, a recessive lethal would show heterozygote advantage if it had a positive pleiotropic effect on production traits thereby conferring an advantage to heterozygote carriers, but not to affected (homozygous) embryos [16]. This would lead to the selection of heterozygote carriers as parents of the next generation, propagate the recessive lethal across population, and in turn significantly affect reproduction [17].

The availability of high-density genotype data has enabled screening of livestock populations for recessive lethals. There are reported successes in dairy cattle [11–14], beef cattle [15], and pigs [18,19]. The screening is based on detecting the absence of homozygotes, either in whole populations or within carrier families. Examples of recessive lethals that also show a positive pleiotropic effect on production include a nonsense mutation in *APAF1* in Holstein cattle [17] and a 660 kbp deletion in Nordic Red cattle [14]. Both of these variations increase milk production compared to wildtype but cause early embryonic death.

Recently, the Irish Cattle Breeding Federation (ICBF) has initiated large-scale genotyping of commercial beef cattle in Ireland [20]. To date more than 1.3 million purebred and crossbred animals have been genotyped. These data will enable genomic selection to increase the productivity and reduce the environmental footprint of Irish cattle. It will also be a powerful resource for biological discovery, including detection of recessive lethals, should they exist.

The objectives of this study were to discover haplotypes carrying putatively recessive lethal alleles in the most numerous Irish beef cattle breeds, and to quantify their pleiotropic effects on traits under selection. We phased the genotypes in sliding windows along the genome for 132,725 purebred animals from five breeds and used five tests to identify haplotypes with absence or reduced homozygosity. We corroborated the identified haplotypes with reproduction and postnatal survival records. We found support for three haplotypes carrying putatively recessive lethal alleles. Substitution analysis showed these haplotypes had pleiotropic effects on economically important traits in Irish beef cattle.

## Materials and methods

In this paper, we analysed several types of data to discover haplotypes carrying putatively recessive lethal alleles in Irish beef cattle. Briefly, the methods consisted of:

(i) Genotype filtering, imputation, and phasing.
(ii) Identification of haplotypes carrying putatively recessive lethal alleles.
(iii) Reproduction and survival analysis to corroborate the identified haplotypes.
(iv) Pleiotropy analysis.
(v) Identification of candidate genes.

### Genotype filtering, imputation, and phasing

We used single nucleotide polymorphism marker array data from the ICBF for purebred Aberdeen Angus, Charolais, Hereford, Limousin, and Simmental animals. The animals were genotyped with four different, but overlapping, Illumina arrays (HD, IDBv1, IDBv2, and IDBv3; Table 1), with most of the animals genotyped on the IDBv3 array (>64%). We filtered the genotype data on autosomal chromosomes, missing information, and heterozygosity. We excluded markers with low genotype call rate (<0.90). We excluded animals with low genotype call rate (<0.90) or high heterozygosity (>6 standard deviations from mean heterozygosity). After filtering we had 132,725 animals: 22,836 Aberdeen Angus, 39,472 Charolais, 12,678 Hereford, 45,965 Limousin, and 11,774 Simmental animals (Table 1). Finally, we reduced the markers to those that were present on the IDBv3 array and had a minor allele frequency above 0.05 within each breed separately. This decreased the number of markers to 40,146 for Aberdeen Angus, 41,366 for Charolais, 40,368 for Hereford, 40,951 for Limousin, and 39,907 for Simmental. We imputed missing genotypes in these reduced sets with AlphaImpute version 1.9.1 [21] and then phased with AlphaPhase version 1.9.1 [21] (using a core and tail length of 320 markers) into 20 marker haplotypes in sliding windows. This means that the genotypes were phased twenty times in sliding windows, with the start of each window being moved along one marker each time.

**Table 1:** Number of markers by array and number of genotyped animals after genotype filtering by breed and array

|  | Array |  |  |  | Total |
| --- | --- | --- | --- | --- | --- |
|  | HD | IDBv1 | IDBv2 | IDBv3 |  |
| Breed / #Markers | 774,959 | 16,044 | 16,291 | 53,368 |  |
| Aberdeen Angus | 532 | 1,969 | 5,409 | 14,926 | 22,836 |
| Charolais | 1,035 | 3,950 | 10,868 | 23,619 | 39,472 |
| Hereford | 362 | 1,759 | 2,825 | 7,732 | 12,678 |
| Limousin | 1,043 | 4,134 | 10,717 | 30,071 | 45,965 |
| Simmental | 417 | 820 | 1,798 | 8,739 | 11,774 |
| Total | 3,389 | 12,632 | 31,617 | 85,087 | 132,725 |

### Identification of haplotypes carrying putatively recessive lethal alleles

To identify haplotypes carrying putatively recessive lethal alleles, the phased haplotypes were analysed for complete absence or reduced level of homozygosity within the analysed breed. We used five different tests that we have grouped into three categories: (i) Population, (ii) Carrier mating, and (iii) Incomplete penetrance. The first and second categories are based on the probability of observing complete absence of recessive homozygous animals in the whole population or in the progeny from carrier matings. Thus, they assume that recessive lethals are completely penetrant. The third category tests the null hypothesis that observed and expected numbers of recessive homozygous animals are equal. Thus, it is able to detect recessive lethals with incomplete penetrance, where some homozygotes do not suffer the deleterious effects or haplotypes imperfectly tag a recessive lethal allele. From this point on, we will use *HH* to denote wildtype homozygotes, *Hh* to denote heterozygotes, and *hh* to denote homozygotes for a haplotype carrying putatively recessive lethal alleles.

#### Population test

The first test identifies haplotypes carrying putatively recessive lethal alleles based on their population frequency. The probability of observing no *hh* animals (*P*_*hh*_) depends on the *Hh* frequency (*C*) and the number of genotyped animals (*N*): *P*_*hh*_ = (1 – *c*^2^/4)^*N*^. The number of expected *hh* animals (*E*_*hh*_) under these conditions is equal to: *E*_*hh*_ = (*N*/4)·*c*^2^. This test is the same as that used by VanRaden et al. [12].

#### Carrier mating tests

The second category consists of two test that identify haplotypes carrying putatively recessive lethal alleles based on the expected number of *hh* progeny from *Hh×Hh* matings. We analysed both sire × dam and sire × maternal grand-sire *Hh* × *Hh* matings. The probability of observing no *hh* animals follows a Bernoulli process and can be calculated as *P*_*hh*_ = 0.75^*C*^ for 6 sire × dam *Hh* × *Hh* matings and *P*_*hh*_ = 0.875^*C*^ for 6 sire × maternal grand-sire *Hh* × *Hh* matings. The expected number of *hh* progeny was calculated as *C*/4 for sire × dam *Hh* × *Hh* matings and *C*/8 for sire × maternal grand-sire *Hh* × *Hh* matings. These tests are partially the same as that used by VanRaden et al. [12].

#### Incomplete penetrance tests

The third category tests the null hypothesis that the observed and expected number of *hh* animals do not differ based on sire × dam *Hh* × *Hh* or sire × maternal grand-sire *Hh* × *Hh* matings. From the number of observed and expected *hh* (*O*_*hh*_ and *E*_*hh*_), *Hh* (*O*_*Hh*_ and *E*_*Hh*_), and *HH* (*O*_*HH*_ and *E*_*HH*_) progeny we calculated a one-way chisquare test statistic: *χ*^2^ = (*O*_*hh*_ − *E*_*hh*_)^2^/*EE*_*hh*_ + (*O*_*HH*+*Hh*_ − *E*_*HH*+*Hh*_)^2^*E*_*HH*+*Hh*_. The test statistic has one degree of freedom.

#### Identification of haplotypes carrying putative recessive lethals

Haplotypes with absence or reduced level of homozygosity at *p* < 5 * 10^-8^ (−*log*_10_*p>* 7.30) were considered as candidates for carrying putatively recessive lethal alleles that would cause early embryonic death or reduced postnatal survival. In order to decrease the false discovery rate only one megabase long regions with at least nine haplotype tests with p < 5 * 10^-8^ were considered. From each of these regions only the most significant haplotype test from each of the three test categories was selected. We analysed the effect of each of these haplotypes on cow reproduction and progeny survival to corroborate lethality and on production traits to evaluate pleiotropy.

### Reproduction and survival analysis to corroborate the identified haplotypes

We used routine cow reproduction and progeny survival records from the ICBF database. We tracked 44,351 insemination records that pertain to purebred Aberdeen Angus, Charolais, Hereford, Limousin, or Simmental cattle, where both the potential sire and the potential dam were genotyped. There were 7,540 records for Aberdeen Angus, 11,520 for Charolais, 5,566 for Hereford, 15,404 for Limousin, and 4,321 for Simmental. Altogether these inseminations resulted in 21,196 progeny. Each resulting calf had records for the date of birth and if deceased the date of death and the cause of death (natural death or slaughtered).

To quantify the effect of identified haplotypes on reproduction, we determined insemination success rate and days between the first insemination and calving. Insemination success rate was calculated as a ratio between the number of born progeny and the number of inseminations. These statistics were reported for the three mating types: (i) *Hh* × *Hh*, (ii) *Hh* × *HH*, and (iii) *HH* × *HH*. The period between the first insemination and calving was calculated from 40,769 records that had known the date of corresponding calving. In order to accommodate shorter or longer than average gestation length a window of time was allowed on either side of the breed average gestation length. We used windows of 7, 14, and 21 days either side of the average breed gestation, calculated as the average gestation length for the set of individuals recorded for that breed. If a calving occurred outside of the window, the insemination was deemed to be unsuccessful. If a haplotype harbours recessive lethals, it is expected that *Hh* × *Hh* matings should have a 25% lower insemination success as there will be ¼ *hh* progeny and more than 21 days longer first insemination to calving interval as this is the average time between each standing oestrus in cattle.

To quantify the effect of identified haplotypes on postnatal survival, we used survival analysis with the Cox proportional-hazard model. We compared hazard ratios between the progeny from: (i) *Hh* × *Hh*, (ii) *Hh* × *HH*, and (iii) *HH* × *HH* matings. If a haplotype affects postnatal survival, we would expect *hh* progeny from *Hh* × *Hh* matings to have higher hazard ratio. Since two causes of death were reported (natural death and slaughtered) we analysed data in two ways. In the first one we treated records from slaughtered animals as censored whereas in the second one we treated their records as uncensored.

### Pleiotropy analysis

We analysed national estimates of breeding values to investigate pleiotropic effect of the identified haplotypes on economically important traits in beef production. Estimated breeding values for 15 traits (feed intake, live weight, cull cow weight, carcass weight, carcass confirmation, carcass fat, docility, gestation length, age at first calving, calving interval, calving difficulty due to the sire, maternal calving difficulty, mortality at calving, maternal weaning weight, and cow survival) for all genotyped individuals in this study were available from the May 2017 routine genetic evaluation carried out by the ICBF. The 15 traits are included in two indexes with different economic weight and trait emphasis (shown in parentheses). The terminal index consists of: calving difficulty due to sire (-€4.65, 18%), gestation length (-€2.25, 4%), mortality (-€5.34, 3%), docility (€17.03, 2%), feed intake (-€0.10, 16%), carcass weight (€3.14, 41%), carcass conformation (€14.77, 11%), and carcass fat (-€7.86, 5%). The replacement index consists of eight cow traits: maternal calving difficulty (-€4.98, 6%), age at first calving (-€0.99, 6%), calving interval (-€5.07, 9%), cow survival (€8.86, 8%), maternal weaning weight (€5.58, 18%), cow live weight (-€1.31, 13%), cow docility (€77.27, 4%), cull cow weight (€0.91, 7%); and eight calf traits: calving difficulty due to sire (-€5.12, 7%), gestation length (-€2.48, 2%), mortality (-€5.87, 1%), calf docility (€14.72, 1%), feed intake (-€0.07, 4%), carcass weight (€2.10, 10%), carcass conformation (€10.22, 3%), and carcass fat (-€5.44, 1%). For every identified haplotype we estimated its substitution effect on estimated breeding values for each of the fifteen traits and two indexes. The substitution effect was estimated by linear regression of estimated breeding values on the number of haplotype copies an animal carries. Trait specific substitution effects were divided with the standard deviation of estimated breeding values to present the effects on a comparable scale.

### Candidate genes

We used Bioconductor version 3.7 with the ENSEMBL_MART_ENSEMBL BioMart database version Ensemble Genes 92 using the btaurus_gene_ensembl to find candidate genes in regions of the identified haplotypes [22]. We used the UMD3.1 cattle reference genome [23]. We extracted the protein coding genes and searched for lethal effects observed in homologous genes with the same function in the Mouse Genome Informatics database [24].

## Results

Our results identify three haplotypes carrying putatively recessive lethal alleles. They were located on chromosome 14 in Aberdeen Angus, chromosome 19 in Charolais, and chromosome 16 in Simmental. The haplotypes show pleiotropic effects on economically important traits in beef production. We identified *ZFAT* as the potential causative gene for lethality in Aberdeen Angus.

### Identification of haplotypes carrying putative recessive lethals

#### Haplotype tests

In total there were between 36 and 109 haplotypes with significant absence or reduced levels of homozygosity at F < 5 * 10^-8^ in the five breeds. Table 2 shows the number of significant haplotypes by test and breed. The highest number of significant haplotypes was observed for Limousin (109) followed by Charolais (86), Aberdeen Angus (69), Hereford (57), and Simmental (36). Of the different tests, the incomplete penetrance test on sire × dam matings had the highest number of significant haplotypes. The carrier mating test on sire × dam matings had the second largest number of significant haplotypes for Charolais, Hereford, and Simmental.

**Table 2:** Number of haplotypes with significant absence or reduced level of homozygosity at F < 5 * 10^-8^ by test and breed

| Test<sup>1</sup> | Aberdeen | Charolais | Hereford | Limousin | Simmental |
| --- | --- | --- | --- | --- | --- |
|  | Angus |  |  |  |  |
| T1 | 3 | 0 | 0 | 8 | 6 |
| T2 | 1 | 9 | 13 | 14 | 15 |
| T3 | 2 | 3 | 0 | 3 | 1 |
| T4 | 65 | 72 | 56 | 90 | 16 |
| T5 | 1 | 5 | 11 | 15 | 1 |
| Total<sup>2</sup> | 69 | 86 | 57 | 109 | 36 |

<sup>1</sup>T1 – Population test; T2 – Carrier mating test on sire × dam matings; T3 – Carrier mating test on sire × maternal grand-sire matings; T4 – Incomplete penetrance test on sire × dam matings; T5 – Incomplete penetrance test on sire × maternal grand-sire matings
<sup>2</sup>Total number of haplotypes with $p < 5 * 10^{-8}$ from at least one test

#### Identification of haplotypes carrying putative recessive lethals

Across all five breeds, there were 19 regions of one megabase in length with at least nine haplotypes with significant absence or reduced level of homozygosity. These were considered as candidate regions with haplotypes carrying putatively recessive lethal alleles. Figure 1 shows the negative logarithm of all test p-values across the whole genome, while Table S1 lists the 19 candidate regions. There were five such regions in Aberdeen Angus, two in Charolais, three in Hereford and Limousin, and six in Simmental. Some of the regions were neighbouring and appear as a single region in the Figure 1. This was the case for three regions on chromosome 14 in Aberdeen Angus, two regions on chromosome 6 in Hereford, two regions on chromosome 23 in Limousin, and three regions on chromosome 13 and two regions on chromosome 16 in Simmental. The region between 48 and 49 megabases on chromosome 19 was discovered in all the breeds.

**Figure 1:**
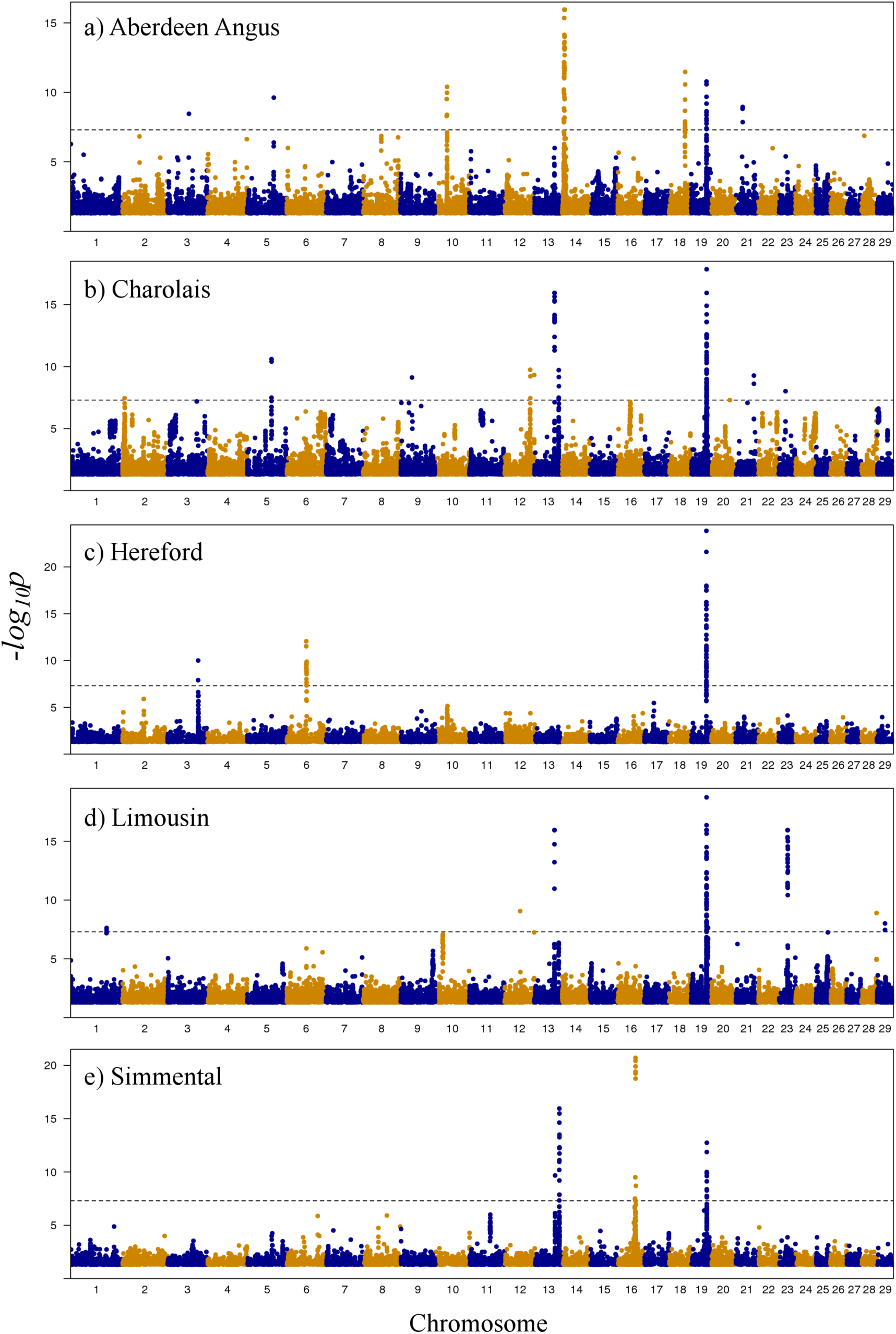
Genome-wide Manhattan plot for the lack of homozygosity in (a) Aberdeen Angus, (b) Charolais, (c) Hereford, (d) Limousin, and (e) Simmental breed.

From the 19 candidate regions, we selected 22 haplotypes with the highest significance within each of the three category tests. Table 3 lists the 22 haplotypes and their characteristics. We named the haplotypes by the breed (AA for Aberdeen Angus, CH for Charolais, HE for Hereford, LI for Limousin and SI for Simmental), the chromosome, and the haplotype number within the breed (H for haplotype). Except for HE19H3 and SI19H6, which were identified with tests from two categories (carrier mating test with sire × dam matings from the carrier mating tests and incomplete penetrance test with sire × dam matings from the incomplete penetrance tests), all haplotypes were identified by tests from a single category.

**Table 3:** Identified haplotypes carrying putatively recessive lethal alleles

| Haplotype name | Test<sup>1</sup> | Chromosome | Haplotype region | Sliding window | Start base | End base | Haplotype percentage |
| --- | --- | --- | --- | --- | --- | --- | --- |
| AA14H1 | T4 | 14 | AA14R1 | 4 | 6,819,966 | 7,514,610 | 25.2 |
| AA14H2 | T4 | 14 | AA14R2 | 9 | 7,646,485 | 8,425,401 | 15.3 |
| AA14H3 | T4 | 14 | AA14R3 | 20 | 8,064,004 | 8,927,881 | 15.2 |
| AA14H4 | T3 | 14 | AA14R3 | 9 | 8,454,690 | 9,215,622 | 15.3 |
| AA18H5 | T4 | 18 | AA14R4 | 4 | 49,446,631 | 50,616,688 | 18.9 |
| AA19H6 | T4 | 19 | AA14R5 | 8 | 48,850,043 | 48,874,999 | 23.9 |
| CH13H1 | T4 | 13 | CH13R1 | 19 | 61,565,261 | 61,584,644 | 33.5 |
| CH19H2 | T4 | 19 | CH19R2 | 5 | 48,874,999 | 48,902,319 | 14.4 |
| CH19H3 | T3 | 19 | CH19R2 | 12 | 48,889,025 | 48,914,435 | 1.2 |
| HE6H1 | T4 | 6 | HE6R1 | 8 | 58,473,731 | 59,706,074 | 30.2 |
| HE6H2 | T4 | 6 | HE6R2 | 4 | 59,688,857 | 60,919,136 | 28.8 |
| HE19H3 | T2, T4 | 19 | HE19R3 | 13 | 48,854,119 | 48,900,428 | 28.9 |
| LI19H1 | T5 | 19 | LI19R1 | 20 | 48,851,566 | 48,871,260 | 12.7 |
| LI19H2 | T2 | 19 | LI19R1 | 11 | 48,883,348 | 48,906,682 | 11.1 |
| LI23H3 | T4 | 23 | LI23R2 | 20 | 27,923,154 | 28,649,349 | 13.7 |
| LI23H4 | T4 | 23 | LI23R3 | 12 | 28,307,706 | 29,872,481 | 11.5 |
| SI13H1 | T2 | 13 | SI13R1, SI13R2 | 15 | 73,746,516 | 74,973,171 | 9.5 |
| SI13H2 | T4 | 13 | SI13R1, SI13R2, SI13R3 | 19 | 73,895,443 | 75,240,419 | 22.3 |
| SI13H3 | T2 | 13 | SI13R2, SI13R3 | 1 | 74,109,992 | 75,310,875 | 9.3 |
| SI16H4 | T2 | 16 | SI16R4, SI16R5 | 16 | 51,195,450 | 53,097,936 | 10.8 |
| SI16H5 | T1 | 16 | SI16R4, SI16R5 | 4 | 51,811,400 | 53,434,159 | 8.8 |
| SI19H6 | T2, T4 | 19 | SI19R6 | 18 | 48,873,060 | 48,891,361 | 18.4 |

<sup>1</sup>T1 – Population test; T2 – Carrier mating test on sire × dam matings; T3 – Carrier mating test on sire × maternal grand-sire matings; T4 – Incomplete penetrance test on sire
× dam matings; T5 – Incomplete penetrance test on sire × maternal grand-sire matings

### Reproduction and survival analysis to corroborate the identified haplotypes

#### Reproduction analysis

Two out of the 22 identified haplotypes, AA14H3 and SI16H5, were associated with decreased insemination success rate and a longer period between the first insemination and calving. Table 4 shows the insemination success rate and the length of the interval between the first insemination and calving for the non-carrier × non-carrier (*HH* × *HH)*, non-carrier × carrier (*HH* × *Hh)*, and carrier × carrier (*Hh* × *Hh)* matings for the AA14H3 and SI16H5 haplotypes. AA14H3 was located on chromosome 14 in Aberdeen Angus between 8,064,004 and 8,927,881 base pairs. SI16A5 was located on chromosome 16 in Simmental between 51,811,400 and 53,434,159 base pairs. The insemination success rates for *Hh* × *Hh* matings were 24.2%, 23.0%, and 20.2% lower (AA14H3), and 48.6%, 52.5%, and 52.7% lower (SI16H5) compared to the *HH* × *HH* matings for a window of 7, 14, and 21 days on either side of the average breed gestation, respectively. The insemination success rates were also lower in *HH* × *Hh* matings than in *HH* × *HH* matings, but the differences were smaller.

**Table 4:**
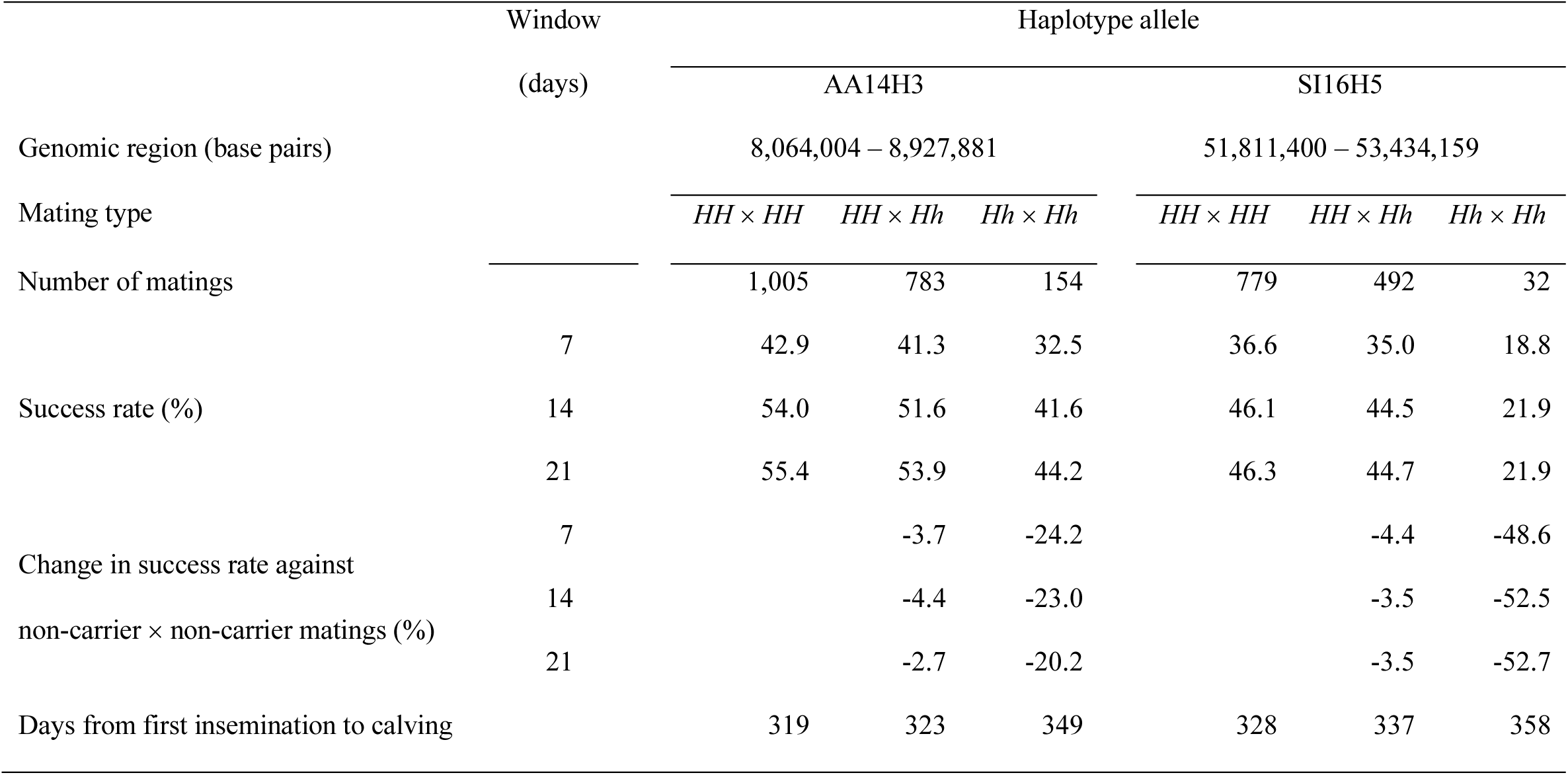
Insemination success rate and interval between the first insemination and calving for non-carrier × non-carrier matings (*HH* × *HH*),non-carrier × carrier (*HH* × *Hh*), and carrier × carrier (*Hh* × *Hh*) matings for AA14H3 and SI16H5 haplotypes and calf registration window of 7,14, and 21 days

|  | Window | Haplotype allele |  |  |  |  |  |
| --- | --- | --- | --- | --- | --- | --- | --- |
|  | (days) | AA14H3 |  |  | SI16H5 |  |  |
| Genomic region (base pairs) |  | 8,064,004 – 8,927,881 |  |  | 51,811,400 – 53,434,159 |  |  |
| Mating type | | $HH \times HH$ | $HH \times Hh$ | $Hh \times Hh$ | $HH \times HH$ | $HH \times Hh$ | $Hh \times Hh$ |
| Number of matings |  | 1,005 | 783 | 154 | 779 | 492 | 32 |
|  | 7 | 42.9 | 41.3 | 32.5 | 36.6 | 35.0 | 18.8 |
| Success rate (%) | 14 | 54.0 | 51.6 | 41.6 | 46.1 | 44.5 | 21.9 |
|  | 21 | 55.4 | 53.9 | 44.2 | 46.3 | 44.7 | 21.9 |
| Change in success rate against | 7 |  | -3.7 | -24.2 |  | -4.4 | -48.6 |
| non-carrier × non-carrier matings (%) | 14 | | -4.4 | -23.0 | | -3.5 | -52.5 |
|  | 21 |  | -2.7 | -20.2 |  | -3.5 | -52.7 |
| Days from first insemination to calving |  | 319 | 323 | 349 | 328 | 337 | 358 |

The period between the first insemination and calving was longer for *Hh* × *Hh* matings than for *HH* × *Hh* matings, which was in turn longer than for *HH* × *HH* matings. The period between the first insemination and calving for *Hh* × *Hh* matings in comparison to *HH* × *HH* matings was longer for 30 days for both AA14H3 and SI16H5 haplotypes. The period between the first insemination to calving for *HH* × *Hh* matings in comparison to *HH* × *HH* matings was longer for 4 days for AA14H3 and 9 days for SI16H5.

#### Survival analysis

There was one identified haplotype, CH19H2, with a significantly (p < 0.05) increased hazard ratio for postnatal survival of progeny from *Hh* × *Hh* matings. CH19H2 was located on chromosome 19 in Charolais between 48,874,999 and 48,902,319 base pairs. Table S2 presents hazard ratios for progeny from *HH* × *Hh* and *Hh* × *Hh* matings as compared to progeny from *HH* × *HH* matings where records for culled progeny were treated as censored or complete. Progeny of individuals that carried CH19H2 had a 36% higher probability of dying or being slaughtered during their life as compared to the progeny of *HH* × *HH* parents.

#### Statistics for haplotypes carrying putatively recessive lethal alleles

The three haplotypes carrying putatively recessive lethal alleles associated with decreased insemination success or reduced postnatal survival all had high frequencies. Table 5 summarizes statistics for the haplotypes AA14A3, CH19A2, and SI16A5. The haplotypes frequencies were 15.2% for AA14A3, 14.4% for CH19A2, and 8.8% for SI16A5. There were 95 recessive homozygous individuals observed for AA14A3, 83 for CH19A2, and none for SI16A5. The table also shows that the number of observed recessive homozygotes was always smaller than the expected number for different types of comparisons. The SNP alleles present on the AA14A3, CH19A2, and SI16A5 haplotypes are shown on Table S3.

**Table 5:** Statistics for the haplotypes AA14H3, CH19H2, and SI16H5

| Haplotype | AA14H3 | CH19H2 | SI16H5 |
| --- | --- | --- | --- |
| Genotyped animals | 22,510 | 38,960 | 11,559 |
| Frequency (%) | 15.2 | 14.4 | 8.8 |
| Recessive homozygotes | 95 | 83 | 0 |
| Expected recessive homozygotes | 194.1 | 256 | 47.6 |
| Sire-carrier × dam-carrier matings | 220 | 256 | 17 |
| Recessive homozygotes from sire-carrier × dam-carrier matings | 2 | 6 | 0 |
| Expected recessive homozygotes from sire-carrier × dam-carrier margins | 55 | 64 | 4.3 |
| Sire-carrier × maternal grand-sire-carrier matings | 141 | 87 | 7 |
| Recessive homozygotes from sire-carrier × maternal grand-sire-carrier matings | 0 | 1 | 0 |
| Expected recessive homozygotes from sire-carrier × maternal grand-sire-carrier matings | 17.6 | 10.9 | 0.9 |
| Probability of not observing recessive homozygotes with the T1 test<sup>1</sup> | $2.3 \cdot 10^{-85}$ | $2.5 \cdot 10^{-112}$ | $2.0 \cdot 10^{-21}$ |
| Probability of not observing recessive homozygotes with the T2 test<sup>1</sup> | $3.3 \cdot 10^{-28}$ | $1.0 \cdot 10^{-32}$ | $7.9 \cdot 10^{-3}$ |
| Probability of not observing recessive homozygotes with the T3 test<sup>1</sup> | $6.7 \cdot 10^{-9}$ | $9.0 \cdot 10^{-9}$ | $3.9 \cdot 10^{-1}$ |
| Probability that the observed and expected number of recessive homozygotes are not different with the T4 test<sup>1</sup> | $1.1 \cdot 10^{-16}$ | $1.1 \cdot 10^{-16}$ | $1.7 \cdot 10^{-2}$ |
| Probability that the observed and expected number of recessive homozygotes are not different with the T5 test<sup>1</sup> | $7.2 \cdot 10^{-6}$ | $1.4 \cdot 10^{-3}$ | $3.2 \cdot 10^{-1}$ |

<sup>1</sup>T1 – Population test; T2 – Carrier mating test on sire × dam matings; T3 – Carrier mating test on sire × maternal grand-sire matings; T4 – Incomplete penetrance test on sire × dam matings; T5 – Incomplete penetrance test on sire × maternal grand-sire mating

### Pleiotropy analysis

To investigate potential pleiotropic effects on traits under selection, we estimated substitution effects of AA14H3, CH19H2, and SI16H5 haplotypes. Figure 2 shows standardized substitution effects for fifteen traits that are part of the terminal or replacement index. A significant substitution effect for AA14H3 haplotype was observed for cull cow weight (0.28), followed by live weight (0.27), carcass weight (0.22), maternal calving difficulty (−0.20), feed intake (0.20), calving interval (0.16), carcass fat (−0.11), calving difficulty (0.10), age at first calving (−0.09), carcass conformation (−0.07), gestation length (0.05), maternal weaning weight (−0.04), and cow survival (0.04). Collectively one copy of the AA14H3 haplotype increases the terminal index for €3.23 where one standard deviation is €18.32 and decreases the replacement index for €3.15, where one standard deviation is €29.52. A significant substitution effect for SI16H5 haplotype was observed for feed intake (−0.19), followed by cull cow weight (0.16), live weight (0.13), carcass conformation (0.11), cow survival (0.11), carcass fat (0.11), and age at first calving (−0.07). Collectively one copy of the SI16H5 haplotype increases the terminal index for €2.30 where one standard deviation is €22.54 and the replacement index for €1.12 where one standard deviation is €35.62. A significant substitution effect for CH19H2 haplotype was observed for maternal weaning weight (−0.16), carcass conformation (0.11), maternal calving difficulty (0.10), carcass fat (−0.10), age at first calving (−0.08), calving difficulty (0.07), live weight (- 0.07), feed intake (−0.07), gestation length (0.07), carcass weight (0.05), and mortality (0.04). Collectively one copy of the CH19H2 haplotype increases the terminal index for €1.47 where one standard deviation is €22.33 and decreases the replacement index for €0.75 where one standard deviation is €30.97.

**Figure 2:**
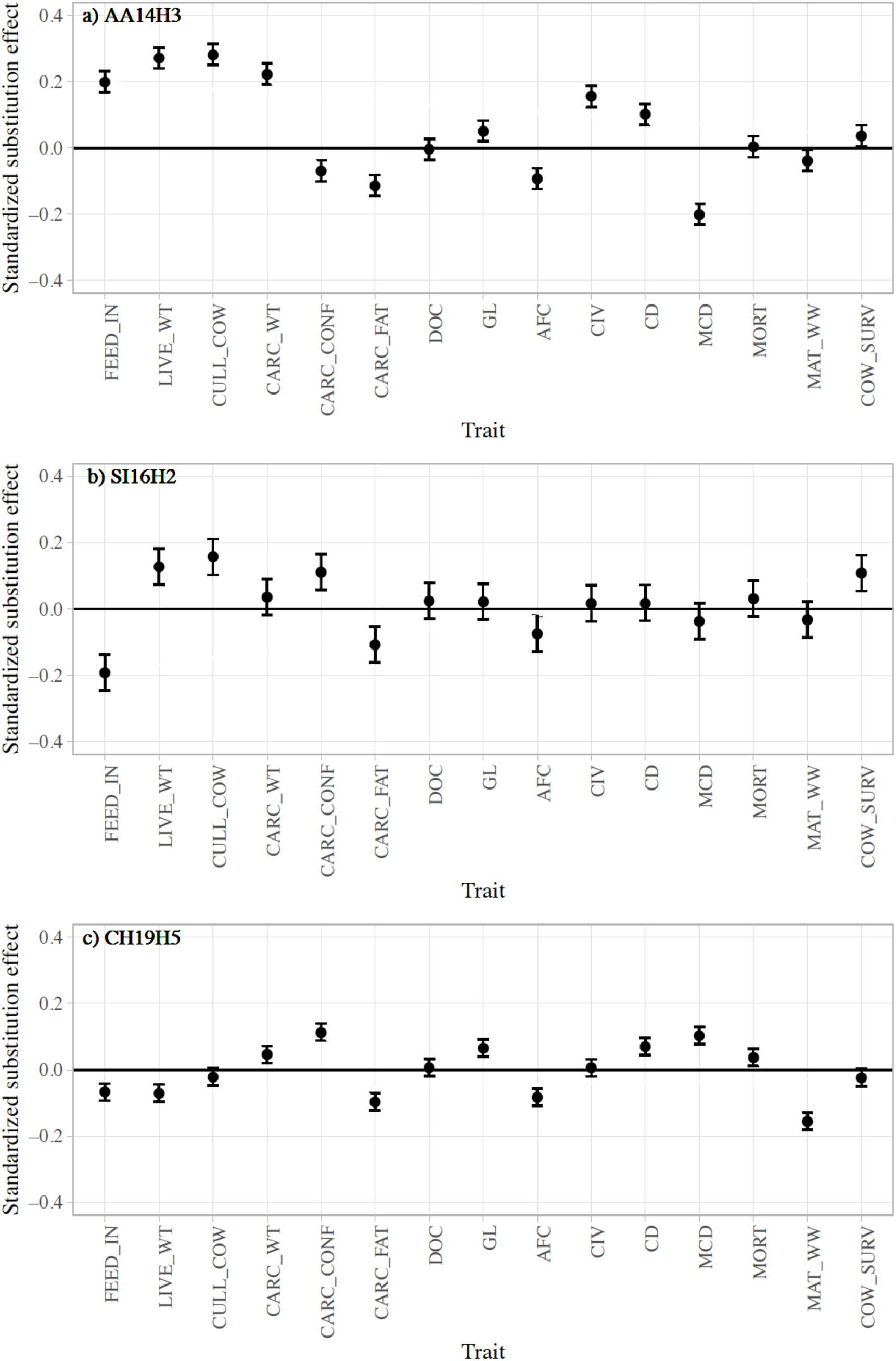
Standardized substitution effects with 95% confidence interval for haplotypes (a) AA14H3, (b) SI16H5, and (c) CH19H2 on 15 traits: FEED_IN – feed intake, LIVE_WT – live weight, CULL_COW – cull cow weight, CARC_WT – carcass weight, CARC_CONF – carcass confirmation, CARC_FAT – carcass fat, DOC – docility, GL – gestation length, AFC – age at first calving, CIV – calving interval, CD – calving difficulty due to the sire, MCD – maternal calving difficulty, MORT – mortality at calving, MAT_WW – maternal weaning weight, COW_SURV– cow survival

### Candidate genes

There was one protein coding gene in the region of AA14H3, none in the region of CH19H2, and several in the region of SI16H5. *Zinc finger and AT-hook domain containing* (*ZFAT*) is the only protein coding gene located between 8,064,004 and 8,927,881 base pairs on chromosome 14 of the AA14A3 haplotype. The *ZFAT* gene is associated with prenatal or perinatal lethality in the Mouse Informatics Database. Table S4 shows protein coding genes located between 51,811,400 and 53,434,159 base pairs on chromosome 16 of the SI16H5 haplotype. There were 64 protein coding genes (Table S4) and five of them (*SKI proto-oncogene, G protein subunit beta 1, ATPase family AAA domain-containing protein 3, and zinc finger and BTB domain containing 17,* caspase-9) are associated with prenatal or perinatal lethality in mice.

## Discussion

The results of this study indicated three haplotypes carrying putatively recessive lethal alleles (AA14A3, CH19A2, and SI16A5) that are candidates for lethality in homozygous state. The three haplotypes were identified by the haplotype analysis of 132,725 purebred animals of five different beef cattle breeds from the ICBF database. The results also suggest pleiotropy, since these haplotypes have substantial substitution effects on economically important traits. In the light of these results, we discuss (i) the identification of haplotypes carrying putative recessive lethals, (ii) the frequency of the identified haplotypes, (iii) their economic impact, and (iv) their potential genetic causes.

### Identification of haplotypes carrying putative recessive lethals

We used five different tests on sliding haplotypes and phenotypic information to search for haplotypes with recessive lethals. We used haplotypes in a sliding window of 20 markers, which was previously shown to be a good compromise between the haplotype diversity and detection of putative recessive lethals with moderate marker density [15]. The tests were based on the number of expected recessive homozygous individuals either in the whole population or from matings between carriers. Along with tests for the complete absence of recessive homozygotes, we also used a set of tests that allowed for some recessive homozygotes but testing whether their number was significantly smaller than expected. This allowed us to take into account possible incomplete penetrance, incomplete linkage between haplotype and putative recessive lethals, structural variation, or genotyping, phasing, and imputation errors. In the following we take all of these to be causes as incomplete penetrance.

The top 22 candidates from the haplotype analysis were further corroborated with cow insemination and progeny survival analysis, and three haplotypes showed phenotypic evidence of lethality. Recessive lethals are expected to decrease insemination success for 25% in carrier (*Hh* × *Hh*) matings, because homozygous embryos do not survive. Further, they are expected to extend the period from the first insemination to calving by 21 days, because this is the average period between two standing oestrus in cattle. In the case of incomplete penetrance some embryos are expected to survive, but might have decreased viability that will in turn affect their survival.

Although there were many haplotypes with significant absence or reduced level of homozygosity, indicating that they carry putatively recessive lethal alleles, only two haplotypes were supported by the reproduction analysis. In particular, there was a strong signal observed at the end of chromosome 19 for all of the five cattle breeds, which was supported with phenotypic data only in Charolais breed. Also, there was a highly significant departure from Hardy-Weinberg equilibrium for markers located at the end of chromosome 19 with excessive heterozygosity (results not shown). This may indicate genotyping errors or structural variation in this region [25], which may explain its strong signal that we observed.

### Frequency of haplotypes carrying putatively recessive lethal alleles

For our methods to detect a haplotype carrying putatively recessive lethal alleles, we needed a relatively high haplotype frequency or many *Hh* × *Hh* matings. Because most haplotypes are rare, this means that the methods we used had sufficient power for a very small proportion of the haplotypes. Following the probability calculations of the different tests and fixing the number of genotyped animals to the number in this study we can assess the minimum required haplotype frequency to reach significance of F < 5 * 10^-8^. The minimum required haplotype frequency to reach significance with the population test in this study was 5.42% for Aberdeen Angus, 4.14% for Charolais, 7.29% for Hereford, 3.83% for Limousin, and 7.56% for Simmental. The percentages of haplotypes with a frequency above these thresholds were very low: 0.98% for Aberdeen Angus, 0.68% for Charolais, 1.01% for Hereford, 0.56% for Limousin, and 0.52% for Simmental. Similarly, reaching significance with the carrier mating test would require at least 59 sire × dam carrier (*Hh* × *Hh*) matings and no homozygous progeny observed. The percentages of haplotypes reaching this threshold were even smaller as for the population test: 0.44% for Aberdeen Angus, 0.23% for Charolais, 0.37% for Hereford, 0.20% for Limousin, and 0.16% for Simmental. Further, reaching significance with the carrier mating test would require 126 sire × maternal grand-sire carrier (*Hh* × *Hh)* matings and no homozygous progeny observed. The percentages of haplotypes reaching this threshold were even smaller as for the carrier mating test with sire × dam matings and were: 0.06% for Aberdeen Angus, 0.04% for Charolais and Hereford, 0.03% for Limousin, and 0.02% for Simmental. Finally, reaching significance with the incomplete penetrance test required chi-square test statistic value higher than 29.72. The percentages of haplotypes that reached this threshold in the current study was <0.01% in all the breeds.

Since recessive lethals are usually rare it is difficult to discover them in a population. This is especially the case for recent mutations since modern breeding programs avoid inbreeding to preserve genetic variation and maintain fitness. Thus, breeders are likely to indirectly avoid carrier matings by virtue of avoiding matings between close relatives. Still, there are some alleles that are propagated through population via popular sires, have pleiotropic effects, or are in incomplete linkage with an allele that has a positive effect on a trait that was selected for in the past. A classic example of a recessive lethal propagated with a popular sire is the complex vertebral malformation (CVM) of calves in Holstein breed. The causing mutation was propagated by the bull Carlin-M Ivanhoe Bell that was incidentally also a carrier of the Bovine leukocyte adhesion deficiency (BLAD) [26]. An example of a pleiotropic recessive lethal is a 660 kbp deletion causing embryonic death in Nordic Red cattle [14]. Since this deletion is positively associated with milk yield, it is quite common (13-32%) in the Nordic Red cattle population. Yet another example is Weaver Syndrome which is expressed postnatally between the age of 5 and 8 months [27]. Here, Weaver-carrier animals also produced more milk and fat, two traits strongly selected in the past [27,28].

High haplotype frequencies and substitution effects on traits under selection suggest that the detected haplotypes in this study may have increased in frequency because of pleiotropy or linked selection. The observed frequencies are high compared to expected frequencies with mutation-selection equilibrium. We can use Nei’s [29] formulas to approximate the expectation and variance of the frequency of a recessive lethal in a finite population, where *μ* is the mutation rate, *N*_*e*_ the effective population size, and *s* the selection coefficient against the mutant lethal:

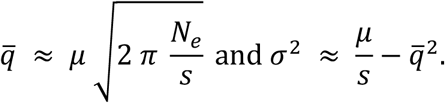

With a high mutation rate of *μ* = 10^-5^ and an effective population size akin to some beef cattle populations of *N*_*e*_ = 300 [e.g., 30,31], the expected equilibrium frequency is very small, regardless of whether the lethal is fully (*s* = 1) or partially penetrant (*s* = 0.75):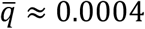 and σ ≈ 0.0031 for a fully penetrant lethal, and 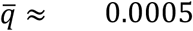 and σ ≈ 0.0036 for a partially penetrant lethal. This analysis clearly shows that there are other forces, i.e., selection, that maintain the high frequency of the three haplotypes.

### Economic impact

Analysis of the economic impact of the three haplotypes on the whole Irish beef population supports the role of pleiotropy in maintaining high haplotype frequencies, which should be managed with appropriately optimised breeding programs. We estimated yearly national economic loss due to early death of homozygous embryos and yearly national economic gain due to improved terminal performance of heterozygous calves. For this purpose we built a simple economic model (see Table S5) based on the number of purebred sire × purebred dam, purebred sire × crossbred dam, and crossbred sire × crossbred dam of Aberdeen Angus, Charolais, and Simmental animals, the expected number of progeny genotypes, haplotype frequencies, insemination success rate, number of days between the subsequent inseminations, daily costs for maintaining a cow, and substitution effects. We obtained these inputs from this study, the ICBF database, and E-profit monitor – an online financial analysis tool [32]. The annual loss was estimated to be about €570,000 for AA14H3, €930,000 for CH19H2, and €210,000 for SI16H5 haplotype. The annual gain was estimated to be about €110,000 for AA14H3, €70,000 for CH19H2 and €50,000 for SI16H5 haplotype. Under the current population structure in Ireland the estimated national annual net effect is negative and is: -€460,000 for AA14H3 -€860,000 for CH19H2, and -€160,000 for SI16H5 haplotype. The large negative net effect for the CH19H2 haplotype is due to the larger purebred population in Charolais than in Aberdeen Angus and Simmental. The economic model shows that under the current population structure the maximum annual net effect would be achieved with haplotype frequencies of 0.02% for AA14H3 (+€6,000), 0.4% for CH19H2 (+€2,000), and 0.02% for SI16H5 (+€4,000) (Figure S1).

### Genetic causes of lethality and pleiotropy

We found one candidate gene for one haplotype carrying putatively recessive lethal alleles, no candidate genes for another, and multiple for the third. *ZFAT* was the only protein coding gene overlapping with the AA14H3 haplotype. *ZFAT* is a transcription factor involved in immune-regulation and apoptosis [33,34] and plays a role in the development of the hematopoietic system [35]. The Mouse Genome Informatics database reported complete early embryonic lethality for a knock-out allele of *ZFAT*. On the other hand *ZFAT* is known to be associated with human height [36–38] and body size in horses [39]. This is consistent with our results, where the insemination success rate was between 20.2% and 24.2% lower (based on different gestation window size) for AA14H3 *Hh* × *Hh* matings as compared to the *HH* × *HH* matings, and the haplotype was positively associated with weight-related traits and feed intake.

There were 64 protein coding genes overlapping with the SI16H5 haplotype. Five of them (*SKI proto-oncogene, G protein subunit beta 1, ATPase family AAA domain-containing protein 3, zinc finger and BTB domain containing 17,* and *caspase-9*) were associated with prenatal or perinatal lethality in the Mouse Genome Informatics database [40–44]. The haplotype has positive association with feed conversion ratio, which may explain its high frequency in the population.

There were no homozygous individuals for the SI16H5 haplotype, which suggests that the identified haplotype tags the putative recessive lethal(s) and that penetrance is complete. The existence of homozygous individuals for AA14H3 and CH19H2 haplotypes suggest there could be multiple underlying haplotypes with and without the lethal(s), but we are unable to distinguish them with current genotype data. Another possible reason is incomplete penetrance, where some homozygotes do not suffer the deleterious effects of the allele.

Fine-mapping these haplotypes with sequence data could potentially resolve the fine haplotype structure, and identify causal variants for lethality and pleiotropic effects on selected traits. Efforts to identify such causative variants are undergoing.

## Conclusions

We identified three haplotypes carrying putatively recessive lethal alleles in purebred Irish beef cattle: one in Aberdeen Angus, one in Charolais, and one in Simmental. Carrier matings in Aberdeen Angus and Simmental had lower success of artificial insemination and longer interval from the first insemination to calving. Progeny from carrier matings in Charolais had higher probability of dying or culling. The haplotype in Aberdeen Angus was in a region that contains the *ZFAT* gene that was previously shown to be lethal in mice. The haplotype in Simmental was in a region that contains to several candidate genes potentially causing prenatal or perinatal lethality. Substitution analysis indicated that the haplotypes had pleiotropic effects on traits under selection. The substitution effects were positive for the terminal index and positive to negative for the replacement index. Further work is needed to identify underlying causative variants impacting traits under selection and fitness. Identification of haplotype carriers will improve the breeding success and fitness of beef cattle populations.

## Authors’ contributions

JJ conceived and designed the study with MM and JMH. GG furthered the ideas and interpretation. JJ performed the analysis. DM prepared phenotypic records. JJ and JMH wrote the first draft. MJ, GG, and JM contributed to interpretation. All the authors read and approved the final manuscript.

### Acknowledgements

Funding for the genotyping of animals used in this study was provided to participants of the Irish Beef Data and Genomics Programme which is a programme funded under Ireland’s Rural Development Programme 2014-2020.

---

**Figure S1:**
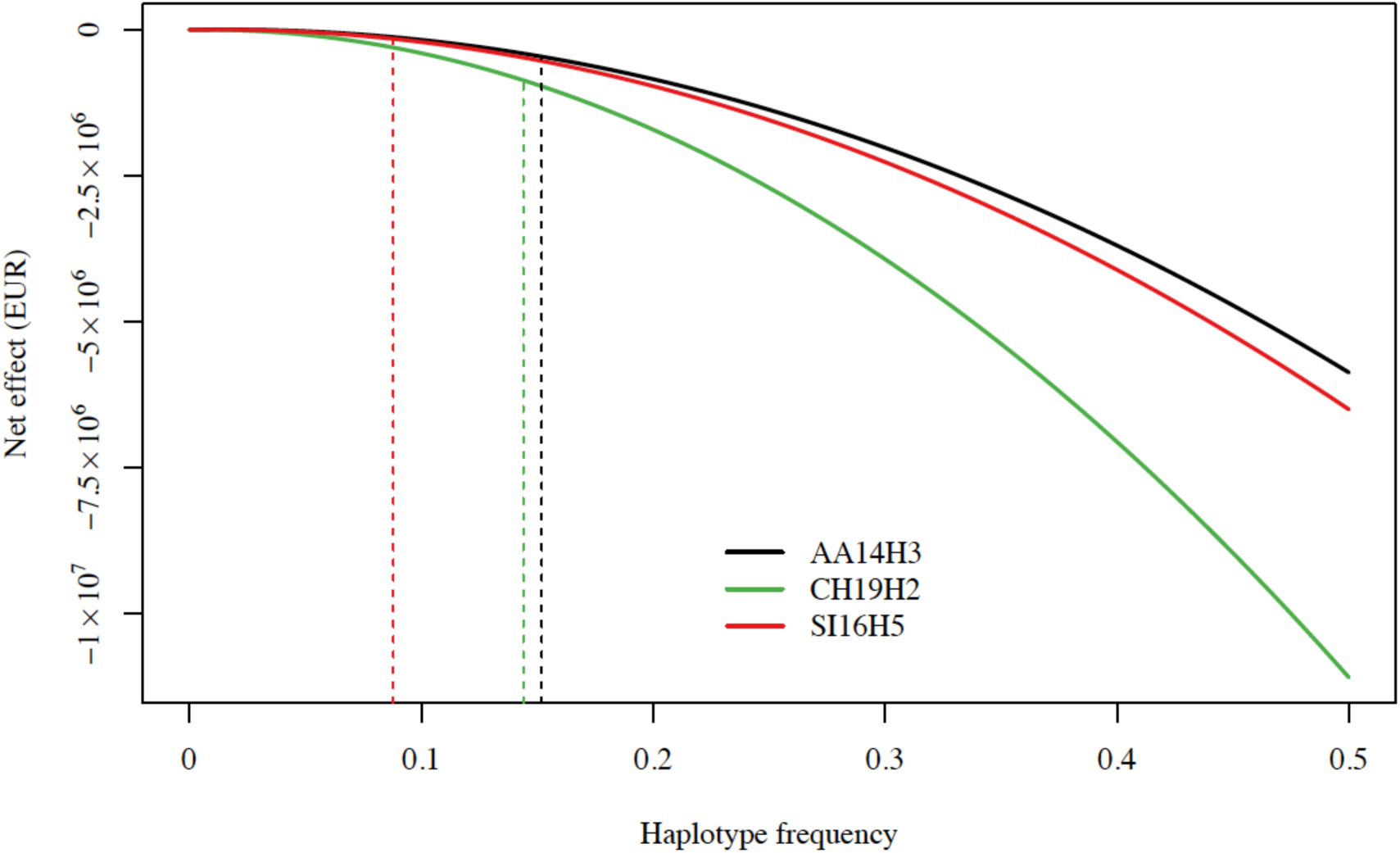
Estimated national annual net effect for the haplotypes AA14H3, CH19H2, and SI16H5 with vertical lines showing the net effect under the current haplotype frequencies

**Table S1:** Genomic regions with at least 9 significant haplotypes for absence or reduced level of homozygosity at *P* < 5 * 10^-8^by breed

| Breed | Region<br>name | Chromosome | Start<br>(Mb) | End<br>(Mb) | Number of<br>haplotypes | Test<br>category<sup>1</sup> |
| --- | --- | --- | --- | --- | --- | --- |
| Aberdeen<br>Angus | AA14R1 | 14 | 6 | 7 | 10 | 3 |
|  | AA14R2 | 14 | 7 | 8 | 24 | 3 |
|  | AA14R3 | 14 | 8 | 9 | 22 | 2, 3 |
|  | AA18R4 | 18 | 49 | 50 | 9 | 3 |
|  | AA19R5 | 19 | 48 | 49 | 12 | 3 |
| Charolais | CH13R1 | 13 | 61 | 70 | 14 | 3 |
|  | CH19R2 | 19 | 48 | 50 | 49 | 2, 3 |
| Hereford | HE6R1 | 6 | 59 | 60 | 16 | 3 |
|  | HE6R2 | 6 | 60 | 61 | 9 | 3 |
|  | HE19R3 | 19 | 48 | 49 | 42 | 2, 3 |
| Limousin | LI19R1 | 19 | 48 | 49 | 50 | 2, 3 |
|  | LI23R2 | 23 | 28 | 29 | 17 | 3 |
|  | LI23R3 | 23 | 29 | 30 | 9 | 3 |
| Simmental | SI13R1 | 13 | 73 | 74 | 9 | 2, 3 |
|  | SI13R2 | 13 | 74 | 75 | 15 | 2, 3 |
|  | SI13R3 | 13 | 75 | 76 | 10 | 2, 3 |
|  | SI16R4 | 16 | 51 | 52 | 9 | 1, 2 |
|  | SI16R5 | 16 | 53 | 54 | 9 | 1, 2 |
|  | SI19R6 | 19 | 48 | 49 | 12 | 2, 3 |

<sup>1</sup>Test category: 1 – Population test, 2 – Carrier mating tests, 3 – Incomplete penetrance tests

**Table S2:** Hazard ratios [95% confidence intervals] for postnatal survival of progeny from non-carrier × carrier (*HH* × *Hh*) and carrier × carrier (*Hh* × *Hh*) matings as compared to the progeny from non-carrier × non-carrier (*HH* × *HH*) matings for the 22 identified haplotypes where

| Haplotype | Censored |  | Complete |  |
| --- | --- | --- | --- | --- |
| | $HH \times Hh$ | $Hh \times Hh$ | $HH \times Hh$ | $Hh \times Hh$ |
| AA14H1 | 1.55 [0.98, 2.46] | 0.77 [0.36, 1.65] | 0.95 [0.77, 1.16] | 0.89 [0.67, 1.17] |
| AA14H2 | 1.70* [1.04, 2.80] | 1.70 [0.74, 3.90] | 0.95 [0.76, 1.19] | 0.92 [0.62, 1.35] |
| AA14H3 | 1.34 [0.79, 2.27] | 2.04 [0.89, 4.70] | 0.86 [0.68, 1.09] | 1.24 [0.84, 1.84] |
| AA14H4 | 1.45 [0.85, 2.47] | 1.28 [0.49, 3.33] | 0.94 [0.75, 1.19] | 0.86 [0.58, 1.28] |
| AA18H5 | 1.43 [0.89, 2.30] | 1.50 [0.78, 2.85] | 0.91 [0.74, 1.12] | 0.99 [0.73, 1.34] |
| AA19H6 | 1.36 [0.83, 2.22] | 1.74 [0.87, 3.44] | 0.80 [0.63, 1.01] | 0.99 [0.66, 1.49] |
| CH13H1 | 0.88 [0.68, 1.14] | 1.05 [0.71, 1.54] | 1.00 [0.87, 1.14] | 1.11 [0.91, 1.36] |
| CH19H2 | 1.00 [0.75, 1.32] | 1.42 [0.86, 2.33] | 1.11 [0.96, 1.28] | 1.36* [1.05, 1.75] |
| CH19H3 | 0.87 [0.64, 1.18] | 0.32 [0.04, 2.26] | 0.93 [0.80, 1.09] | 0.67 [0.35, 1.29] |
| HE6H1 | 0.93 [0.62, 1.40] | 0.79 [0.42, 1.49] | 1.24* [1.02, 1.50] | 1.01 [0.75, 1.36] |
| HE6H2 | 0.84 [0.57, 1.26] | 0.89 [0.52, 1.51] | 1.17 [0.97, 1.41] | 1.10 [0.86, 1.41] |
| HE19H3 | 1.37 [0.92, 2.03] | 1.17 [0.58, 2.34] | 1.21 [0.98, 1.49] | 1.16 [0.73, 1.83] |
| LI19H1 | 1.10 [0.87, 1.40] | 1.03 [0.63, 1.68] | 0.90 [0.80, 1.01] | 1.04 [0.82, 1.31] |
| LI19H2 | 0.97 [0.72, 1.31] | 0.66 [0.21, 2.06] | 0.97 [0.84, 1.11] | 0.63 [0.35, 1.15] |
| LI23H3 | 0.94 [0.73, 1.23] | 0.85 [0.47, 1.52] | 0.92 [0.82, 1.04] | 0.84 [0.65, 1.09] |
| LI23H4 | 1.02 [0.78, 1.33] | 1.47 [0.86, 2.49] | 0.83* [0.73, 0.95] | 0.94 [0.71, 1.26] |
| SI13H1 | 0.99 [0.53, 1.87] | 1.50 [0.45, 4.94] | 0.98 [0.75, 1.28] | 1.21 [0.67, 2.17] |
| SI13H2 | 0.88 [0.54, 1.46] | 1.60 [0.89, 2.89] | 0.95 [0.78, 1.17] | 1.30 [0.99, 1.71] |
| SI13H3 | 0.84 [0.43, 1.67] | 1.85 [0.65, 5.29] | 0.94 [0.71, 1.23] | 1.05 [0.59, 1.85] |
| SI16H4 | 1.09 [0.64, 1.86] | 1.06 [0.32, 3.50] | 1.08 [0.87, 1.35] | 1.09 [0.67, 1.78] |
| SI16H5 | 1.07 [0.59, 1.97] | 2.57 [0.35, 18.95] | 1.25 [0.98, 1.60] | 2.36 [0.87, 6.40] |
| SI19H6 | 1.06 [0.62, 1.83] | 1.04 [0.40, 2.70] | 1.02 [0.80, 1.29] | 1.17 [0.80, 1.71] |

\*p < 0.05

**Table S3:**
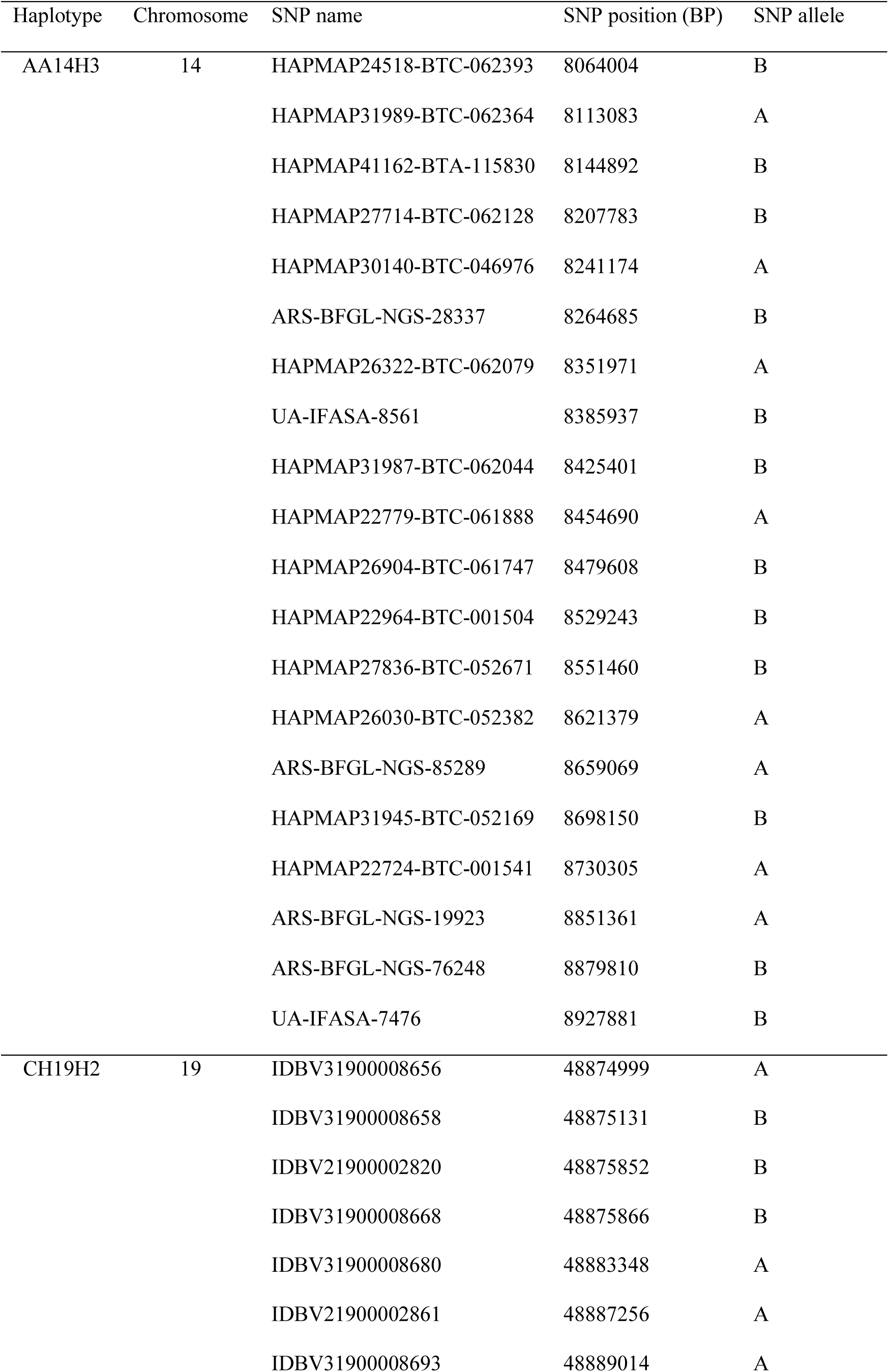
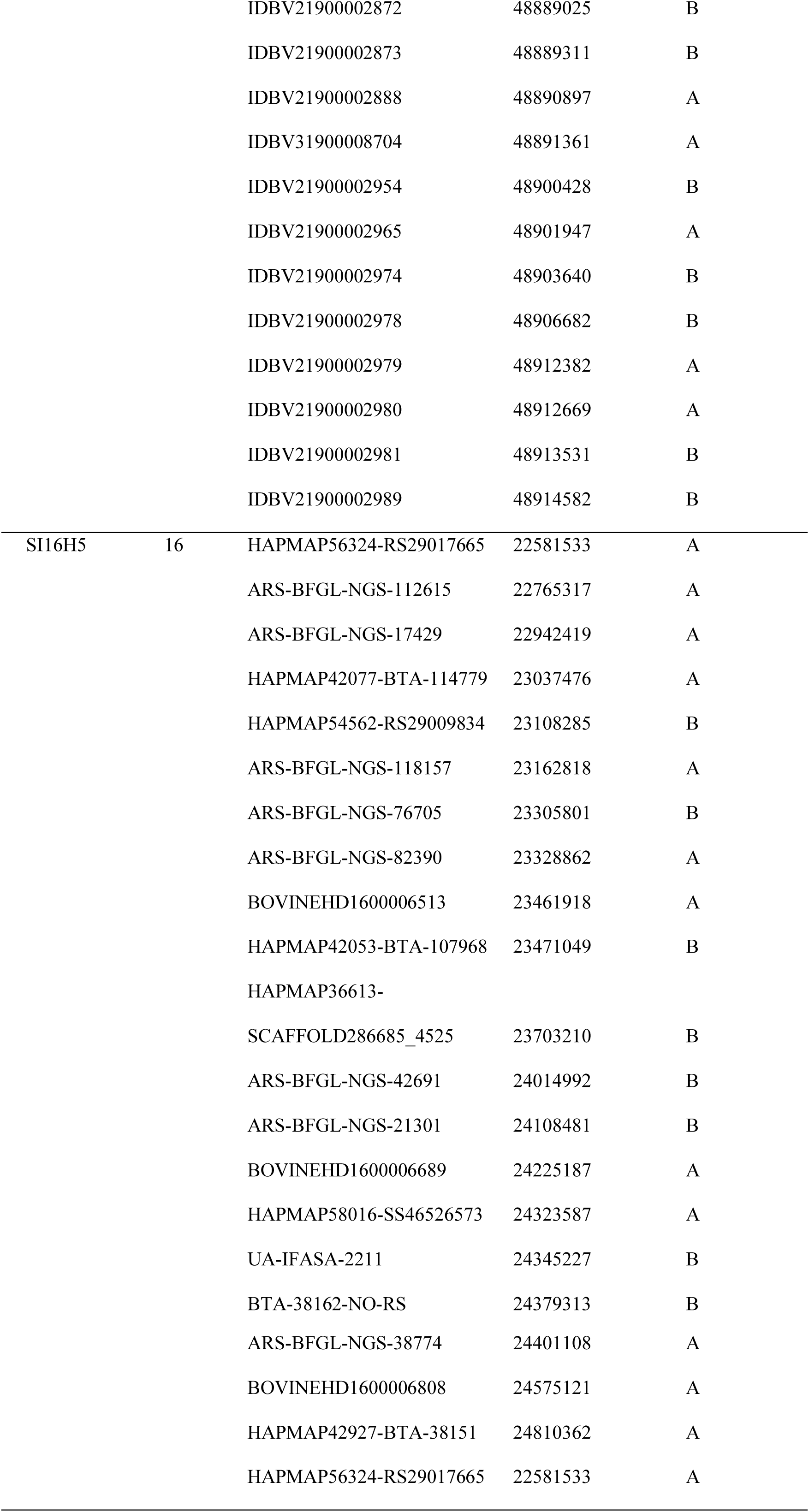
SNP alleles on the AA14H3, CH19H2, and SI16H5 haplotype

**Table S4:**
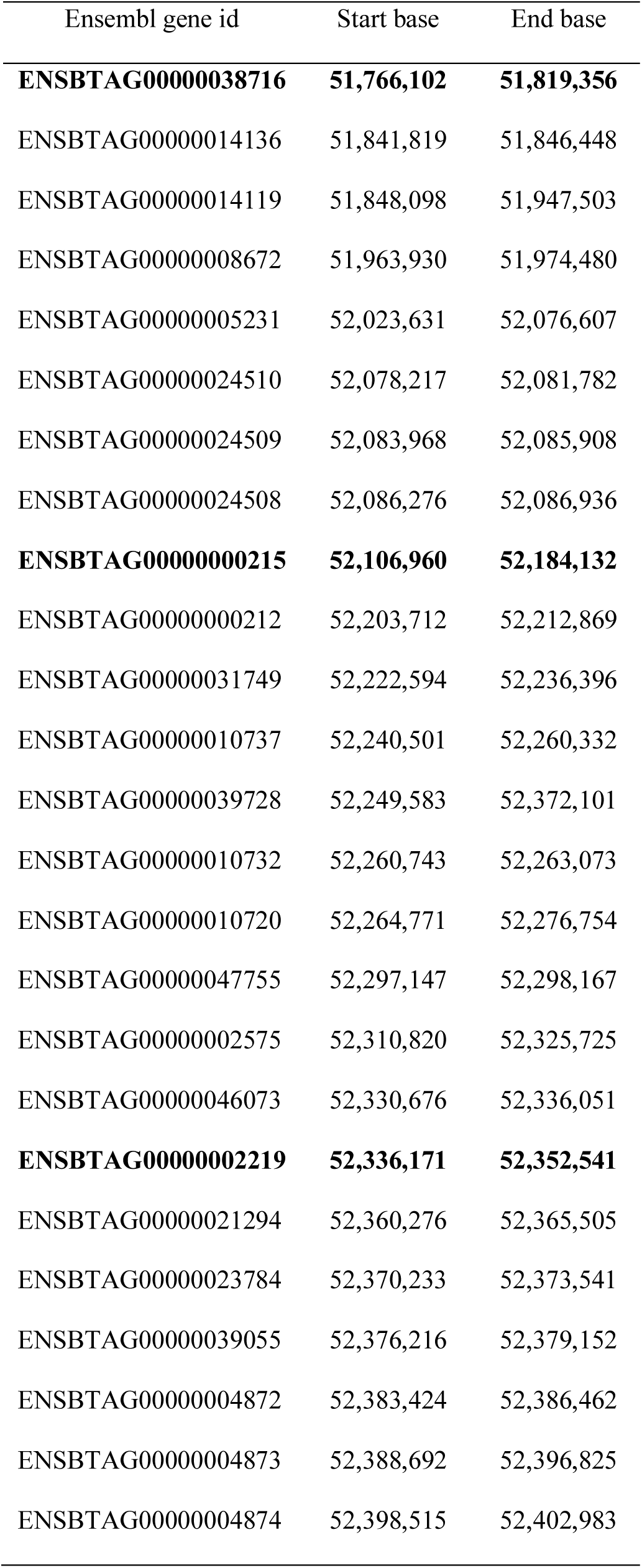
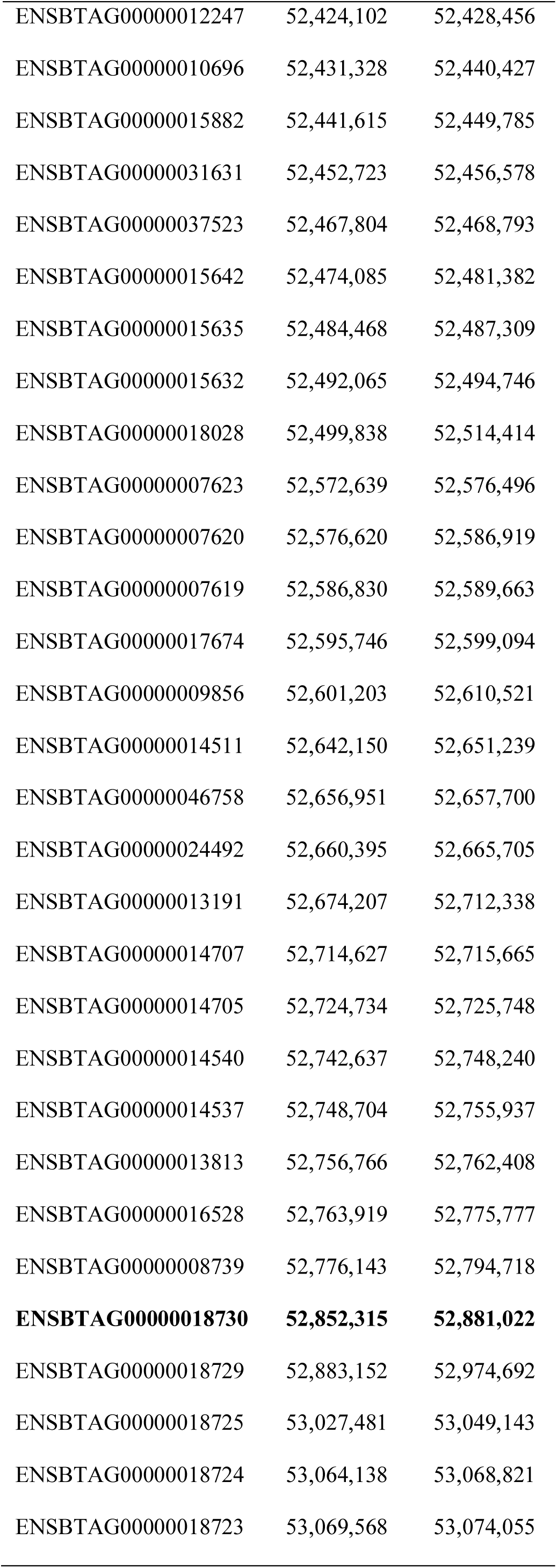
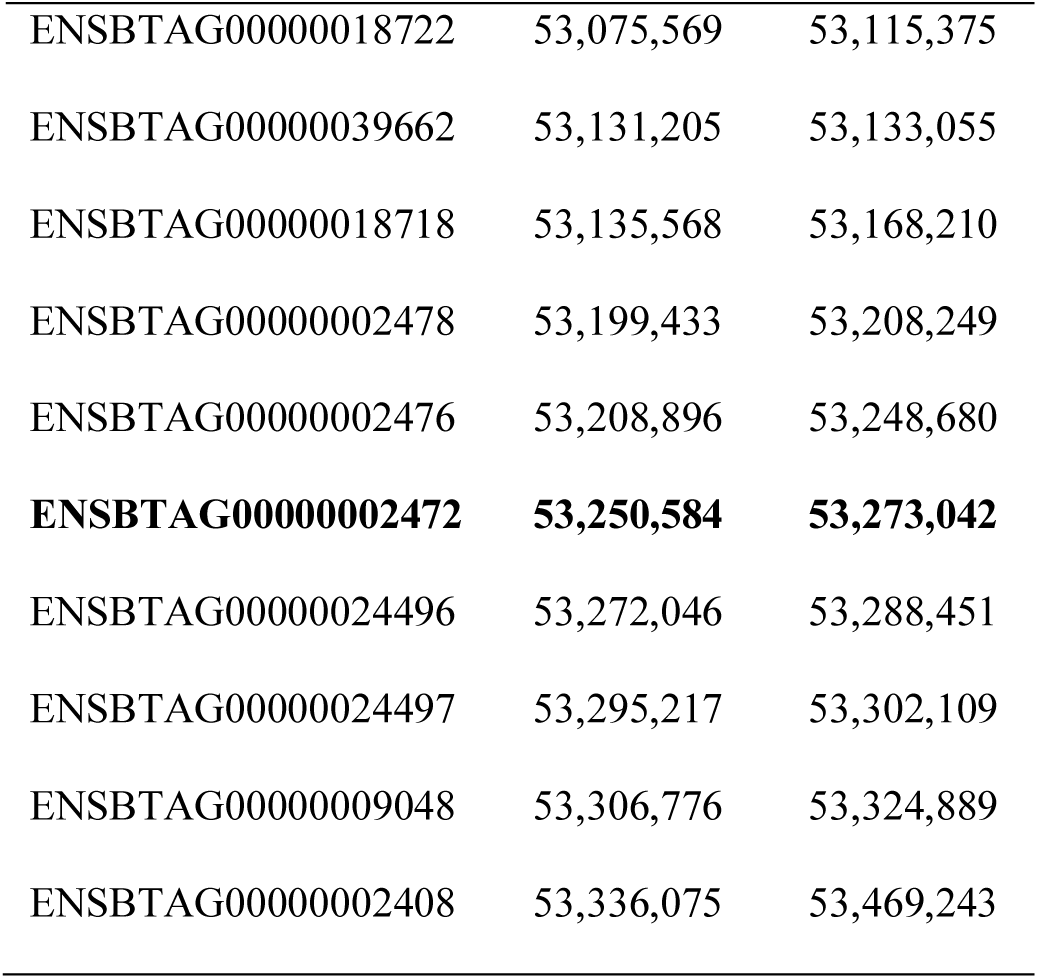
Protein coding genes between 51,611,400 and 53,234,159 base pairs of bovine chromosome 16 of the SI16H5 haplotype with genes showing prenatal or perinatal lethality in mice in bold

| Ensembl gene id | Start base | End base |
| --- | --- | --- |
| **ENSBTAG00000038716** | **51,766,102** | **51,819,356** |
| ENSBTAG00000014136 | 51,841,819 | 51,846,448 |
| ENSBTAG00000014119 | 51,848,098 | 51,947,503 |
| ENSBTAG00000008672 | 51,963,930 | 51,974,480 |
| ENSBTAG00000005231 | 52,023,631 | 52,076,607 |
| ENSBTAG00000024510 | 52,078,217 | 52,081,782 |
| ENSBTAG00000024509 | 52,083,968 | 52,085,908 |
| ENSBTAG00000024508 | 52,086,276 | 52,086,936 |
| **ENSBTAG00000000215** | **52,106,960** | **52,184,132** |
| ENSBTAG00000000212 | 52,203,712 | 52,212,869 |
| ENSBTAG00000031749 | 52,222,594 | 52,236,396 |
| ENSBTAG00000010737 | 52,240,501 | 52,260,332 |
| ENSBTAG00000039728 | 52,249,583 | 52,372,101 |
| ENSBTAG00000010732 | 52,260,743 | 52,263,073 |
| ENSBTAG00000010720 | 52,264,771 | 52,276,754 |
| ENSBTAG00000047755 | 52,297,147 | 52,298,167 |
| ENSBTAG00000002575 | 52,310,820 | 52,325,725 |
| ENSBTAG00000046073 | 52,330,676 | 52,336,051 |
| **ENSBTAG00000002219** | **52,336,171** | **52,352,541** |
| ENSBTAG00000021294 | 52,360,276 | 52,365,505 |
| ENSBTAG00000023784 | 52,370,233 | 52,373,541 |
| ENSBTAG00000039055 | 52,376,216 | 52,379,152 |
| ENSBTAG00000004872 | 52,383,424 | 52,386,462 |
| ENSBTAG00000004873 | 52,388,692 | 52,396,825 |
| ENSBTAG00000004874 | 52,398,515 | 52,402,983 |
| ENSBTAG00000012247 | 52,424,102 | 52,428,456 |
| ENSBTAG00000010696 | 52,431,328 | 52,440,427 |
| ENSBTAG00000015882 | 52,441,615 | 52,449,785 |
| ENSBTAG00000031631 | 52,452,723 | 52,456,578 |
| ENSBTAG00000037523 | 52,467,804 | 52,468,793 |
| ENSBTAG00000015642 | 52,474,085 | 52,481,382 |
| ENSBTAG00000015635 | 52,484,468 | 52,487,309 |
| ENSBTAG00000015632 | 52,492,065 | 52,494,746 |
| ENSBTAG00000018028 | 52,499,838 | 52,514,414 |
| ENSBTAG00000007623 | 52,572,639 | 52,576,496 |
| ENSBTAG00000007620 | 52,576,620 | 52,586,919 |
| ENSBTAG00000007619 | 52,586,830 | 52,589,663 |
| ENSBTAG00000017674 | 52,595,746 | 52,599,094 |
| ENSBTAG00000009856 | 52,601,203 | 52,610,521 |
| ENSBTAG00000014511 | 52,642,150 | 52,651,239 |
| ENSBTAG00000046758 | 52,656,951 | 52,657,700 |
| ENSBTAG00000024492 | 52,660,395 | 52,665,705 |
| ENSBTAG00000013191 | 52,674,207 | 52,712,338 |
| ENSBTAG00000014707 | 52,714,627 | 52,715,665 |
| ENSBTAG00000014705 | 52,724,734 | 52,725,748 |
| ENSBTAG00000014540 | 52,742,637 | 52,748,240 |
| ENSBTAG00000014537 | 52,748,704 | 52,755,937 |
| ENSBTAG00000013813 | 52,756,766 | 52,762,408 |
| ENSBTAG00000016528 | 52,763,919 | 52,775,777 |
| ENSBTAG00000008739 | 52,776,143 | 52,794,718 |
| **ENSBTAG00000018730** | **52,852,315** | **52,881,022** |
| ENSBTAG00000018729 | 52,883,152 | 52,974,692 |
| ENSBTAG00000018725 | 53,027,481 | 53,049,143 |
| ENSBTAG00000018724 | 53,064,138 | 53,068,821 |
| ENSBTAG00000018723 | 53,069,568 | 53,074,055 |
| ENSBTAG00000018722 | 53,075,569 | 53,115,375 |
| ENSBTAG00000039662 | 53,131,205 | 53,133,055 |
| ENSBTAG00000018718 | 53,135,568 | 53,168,210 |
| ENSBTAG00000002478 | 53,199,433 | 53,208,249 |
| ENSBTAG00000002476 | 53,208,896 | 53,248,680 |
| **ENSBTAG00000002472** | **53,250,584** | **53,273,042** |
| ENSBTAG00000024496 | 53,272,046 | 53,288,451 |
| ENSBTAG00000024497 | 53,295,217 | 53,302,109 |
| ENSBTAG00000009048 | 53,306,776 | 53,324,889 |
| ENSBTAG00000002408 | 53,336,075 | 53,469,243 |

**Table S5:** Estimated economic effect for the haplotypes AA14H3, CH19H2, and SI16H5

| Haplotype | AA14H3 | CH19H2 | SI16H5 |
| --- | --- | --- | --- |
| Number of calves with purebred sire and purebred dam | 9,898 | 12,100 | 4,306 |
| Number of calves with purebred sire and crossbred dam | 136,081 | 218,893 | 170,648 |
| Number of calves with crossbred sire and crossbred dam | 19,277 | 16,230 | 9,037 |
| Haplotype frequency | 0.152 | 0.144 | 0.088 |
| Insemination success rate | 0.520 | 0.423 | 0.452 |
| Number of days between two inseminations | 70 | 69 | 60 |
| Haplotype substitution effect for terminal index | 3.227 | 1.469 | 2.298 |
| Costs of keeping cow in herd (€) | 2.2 | 2.2 | 2.2 |
| Number of homozygous embryos | 3,677 | 6,157 | 1,574 |
| Economic loss due to the lethality (€) | 566,257 | 934,673 | 207,810 |
| Number of heterozygous calves born | 33,141 | 47,894 | 22,655 |
| Economic gain for terminal index (€) | 106,946 | 70,356 | 52,061 |
| Net effect (€) | -459,311 | -864,317 | -155,748 |

